# Evolutionarily new genes in humans with disease phenotypes reveal functional enrichment patterns shaped by adaptive innovation and sexual selection

**DOI:** 10.1101/2023.11.14.567139

**Authors:** Jian-Hai Chen, Patrick Landback, Deanna Arsala, Alexander Guzzetta, Shengqian Xia, Jared Atlas, Dylan Sosa, Yong E. Zhang, Jingqiu Cheng, Bairong Shen, Manyuan Long

**Affiliations:** Department of Ecology and Evolution, The University of Chicago, 1101 E 57th Street, Chicago, IL 60637; Institutes for Systems Genetics, West China University Hospital, Chengdu 610041, China; Department of Pathology, The University of Chicago, 1101 E 57th Street, Chicago, IL 60637; Key Laboratory of Zoological Systematics and Evolution, Institute of Zoology, Chinese Academy of Sciences, Beijing 100101, China

**Keywords:** New genes, Pleiotropy, Young genes, Phenotypic innovation, Sexual selection, Natural selection

## Abstract

New genes (or young genes) are genetic novelties pivotal in mammalian evolution. However, their phenotypic impacts and evolutionary patterns over time remain elusive in humans due to the technical and ethical complexities of functional studies. Integrating gene age dating with Mendelian disease phenotyping, our research shows a gradual rise in disease gene proportion as gene age increases. Logistic regression modeling indicates that this increase in older genes may be related to their longer sequence lengths and higher burdens of deleterious de novo germline variants (DNVs). We also find a steady integration of new genes with biomedical phenotypes into the human genome over macroevolutionary timescales (∼0.07% per million years). Despite this stable pace, we observe distinct patterns in phenotypic enrichment, pleiotropy, and selective pressures across gene ages. Notably, young genes show significant enrichment in diseases related to the male reproductive system, indicating strong sexual selection. Young genes also exhibit disease-related functions in tissues and systems potentially linked to human phenotypic innovations, such as increased brain size, musculoskeletal phenotypes, and color vision. We further reveal a logistic growth pattern of pleiotropy over evolutionary time, indicating a diminishing marginal growth of new functions for older genes due to intensifying selective constraints over time. We propose a “pleiotropy-barrier” model that delineates higher potentials for phenotypic innovation in young genes compared to older genes, a process that is subject to natural selection. Our study demonstrates that evolutionarily new genes are critical in influencing human reproductive evolution and adaptive phenotypic innovations driven by sexual and natural selection, with low pleiotropy as a selective advantage.

## Introduction

The imperfection of DNA replication serves as a rich source of variations for evolution and biodiversity (Nei 2013). Such genetic variations underpin the ongoing evolution of human phenotypes, with beneficial mutations being fixed by positive selection, and detrimental ones being eliminated through purifying selection. In medical terminology, this spectrum is categorized as "case and control" or "disease and health," representing two ends of the phenotypic continuum (Pavličev and Wagner 2022). Approximately 8,000 clinical types of rare Mendelian disorders, affecting millions worldwide, are attributed to deleterious DNA mutations in single genes (monogenic) or a small number of genes (oligogenic) with significant effects (Antonarakis and Beckmann 2006; Fetro and Scherman 2020). To date, over 4,000 Mendelian disease genes have been identified, each contributing to a diverse array of human phenotypes (https://mirror.omim.org/statistics/geneMap) (Boycott et al. 2013). These identified disease genes and associated phenotypes could provide critical insights into the evolutionary trajectory of human traits (Claussnitzer et al. 2020).

Evolutionarily new genes – such as de novo genes, chimeric genes, and gene duplicates – have been continually emerging and integrating into the human genome throughout microevolutionary processes (Brosius 1991; Kaessmann et al. 2002; Conrad and Antonarakis 2007; Baertsch et al. 2008; Wu et al. 2011; Long et al. 2013; Van Oss and Carvunis 2019; Betrán and Long 2022; Zhang et al. 2022). New genes can be integrated into biologically critical processes, such as transcription regulation, RNA synthesis, and DNA repair (Ciccarelli et al. 2005; Ding et al. 2021). For instance, in yeast, some new genes may play roles in DNA repair processes (Cai et al. 2008; Li et al. 2010; Parikh et al. 2022). In *Drosophila* species, lineage-specific genes may control the key cytological process of mitosis (Ross et al. 2013). New genes have also been found with roles in early larval development of *Drosophila* (Kasinathan et al. 2020). In nematodes, insects, and fish, some lineage-specific genes are thought to be involved in morphological development, a process which was long believed to be governed by deeply conserved genetic mechanisms (Ragsdale et al. 2013; Klomp et al. 2015; Li et al. 2021). These studies from model species reveal various important biological functions of new genes.

Compared to non-human model organisms, where gene functions can be characterized through genetic knockdowns and knockouts, investigating functions of human genes in their native context is impractical. Despite this limitation, accumulating omics data and *in vitro* studies of human genes have suggested the potential roles of evolutionarily young genes in basic cellular processes and complex phenotypic innovations (Marques et al. 2005; Shi and Su 2012; Zhang and Long 2014). Brain transcriptomic analysis has revealed that up-regulated genes early in human development are enriched with primate-specific genes, particularly within the human-specific prefrontal cortex (Zhang et al. 2011). The recruitment of new genes into the transcriptome suggests that human anatomical novelties may evolve with the contribution of new gene evolution. Growing evidence from recent decades regarding the functions of new genes contradicts the conventional conservation-dominant view of human genetics and phenotypes.

It has long been observed that there are more disease genes among older genes than among young ones (Domazet-Lošo and Tautz 2008). However, the underlying mechanism remains unclear. In recent years, the increasing availability of large-scale sequences of whole exomes (WES) and whole genomes (WGS) has identified deleterious variants causing rare disorders, “orphan” diseases, and rare forms of common diseases (Richards et al. 2015). Rare diseases are often caused by rare variants, which have greater effects on disease phenotypes than common variants (Richards et al. 2015; Wang et al. 2021; Chen et al. 2022a; Greene et al. 2023; Jia et al. 2023; Weiner et al. 2023; Chen et al. 2024a). The effect of gene-based rare variant burden—the aggregate impact of rare (including de novo germline) genetic variants in cohorts—has been confirmed in many genetic disorders (Purcell et al. 2014; Zuk et al. 2014; Guo et al. 2018; Halvorsen et al. 2020; Jiang et al. 2021).

In this study, we analyzed the anatomical organ/tissue/system phenotypes (OP) of human genetic diseases to understand questions about gene ages, phenotypic enrichment, pleiotropy, and selective constraints. First, are older genes more likely to be disease genes, and if so, why? Second, does the rate of disease gene emergence (per million years) differ across macroevolutionary history? Third, do young genes show a phenotypic preference for certain disease systems? Fourth, do young genes exhibit a different pattern of pleiotropic effects compared to older genes? Finally, are these differences driven by selection?

## Results

### Young genes have lower fractions of disease genes with organs/tissues phenotypes than older genes

We determined the evolutionary ages (phylostrata) for 19,665 genes shared between the GenTree database (Shao et al. 2019) and Ensembl (v110) (Supplemental Table S1). These genes were then categorized into two types: “disease genes” or “non-disease genes”, based on disease annotations for a total of 5006 genes from the Human Phenotype Ontology database (HPO, September 2023), which is the de facto standard for phenotyping rare Mendelian diseases (Köhler et al. 2018). Among these disease genes, 60 genes have no gene age information. Thus, an intersection of datasets yielded 4,946 genes annotated with both evolutionary age and organ/tissue/system-specific phenotypic abnormalities (Fig. 1A-1B and Supplemental Table 2). To ensure sufficient statistical power in comparisons, we merged evolutionary age groups with a small number of genes (<100) with their adjacent older group (Fig. 1A). As a result, we reclassified these genes into seven ancestral age groups, ranging from Euteleostomi (and more ancient) nodes to modern humans (br0-br6, Fig. 1A). We observed an increase in the proportion of disease genes over evolutionary time (Supplemental Fig. S1,Figs. 1A and 1C), suggesting that gene age impacts disease susceptibility—a trend qualitatively consistent with earlier studies (Domazet-Lošo and Tautz 2008).

**Figure 1.**
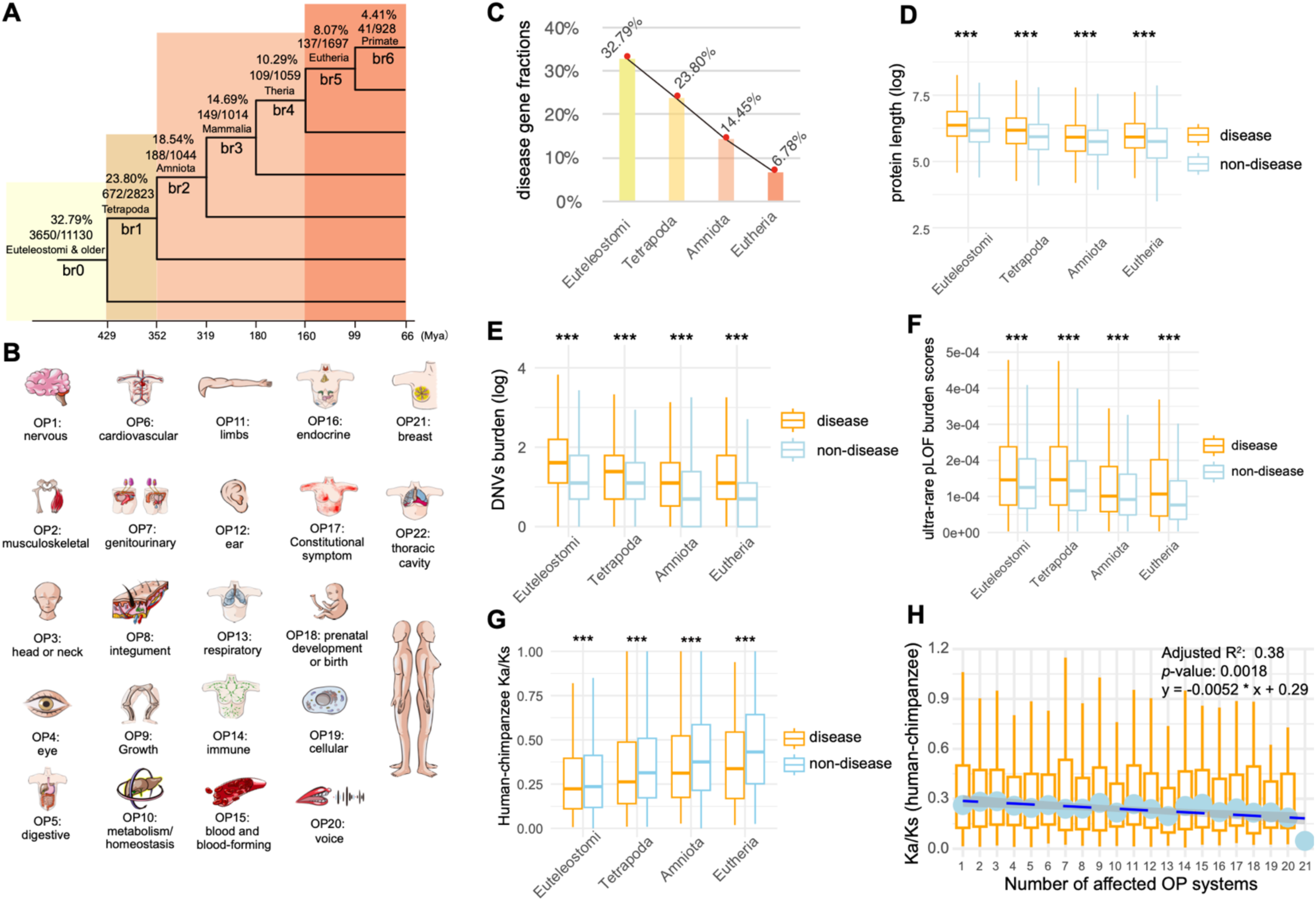
Number of genes, burdens of deleterious rare variants, and Ka/Ks ratios for genes categorized by gene ages (phylostrata) and disease systems (OP). (*A*) The phylogenetic framework illustrating phylostrata and disease genes associated with disease systems (OP). The phylogenetic branches represent phylostratum assignment for all genes and disease genes. The "br" values from br0 to br6 indicate different age groups (or branches). These are further categorized into four phylostrata to resolve the small number problem in “br6” for some analyses. The horizontal axis blow depicts the divergence time sourced from the Timetree database (July 2023). The numbers of total genes and disease genes and their ratios are shown for each phylostratum. (*B*) The 22 HPO-defined organ/tissue systems (OP), which are ordered based on the fraction of disease genes in certain system out of all disease genes. (*C*) The fractions of disease genes at four major phylostrata from Euteleostomi to Eutheria. (*D*) The protein lengths between disease and non-disease genes across four phylostrata. (*E*) The gene-wise burden based on de novo germline variants from Gene4Denovo database (Zhao et al. 2020) between disease and non-disease genes across four phylostrata. (*F*) The burden score of ultrarare predicted loss-of-function (pLOF) variants (Weiner et al. 2023) between disease and non-disease genes across four phylostrata. (*G*) The pairwise Ka/Ks ratios from the Ensembl database based on Maximum Likelihood estimation for “one-to-one” orthologs between human and chimpanzee. Only genes under purifying selection are shown (Ka/Ks < 1, 3226 genes). (*H*) The boxplot for the number of affected systems for disease genes and their pairwise Ka/Ks ratios for “one-to-one” orthologs between human and chimpanzee (Ka/Ks ratios for all 3369 genes). The linear regression is based on median values and the number of affected tissues, with the statistical details and formula displayed in the upper right corner. Note: all significance levels of comparisons between disease genes and non-disease genes are determined using the Wilcoxon rank sum test. The symbol “***” indicates significance level of *p* < 0.001.

### The likelihood of being disease genes positively correlates with gene age, protein length, and DNVs burden

To quantify factors contributing to the likelihood of a gene being classified as a disease gene, we assigned binary states to all genes (“1”, disease genes; “0”, non-disease genes) and performed stratified logistic regression modeling (Supplemental Table S4). We explored multiple predictors, including gene age (*T*, mya), gene length (*L_g_*) or protein length (*L*), gene-wise burden of deleterious de novo germline mutations (DNVs, *D*) from 46,612 trios (Wang et al. 2022), and rare variant burden (*R*) based on gnomAD genomes (Chen et al. 2024b) (Supplemental Table S3). Sequence lengths were considered because, assuming a random mutation process, longer sequences would be expected to accumulate more deleterious variants in a cohort. Additionally, metrics of rare variant burdens were considered because Mendelian disorders and rare forms of common diseases are predominantly influenced by rare variants due to their significant phenotypic effects (Gibson 2012; Guo et al. 2018; Kingdom et al. 2024). The burden of rare variants, which is often based on a collapsed information of rare loss-of-function or deleterious variants within causal genes at the cohort level, has been actively utilized in studying rare diseases (Lee et al. 2014).

Here, the burden scores of rare variants were based on two datasets: gene-wise burden of predicted deleterious de novo germline mutations (DNVs) from 46,612 trios (Wang et al. 2022) and gene-wise burden of rare variants from 76,215 genomes in the gnomAD database (v4.0.0, minor allele frequency (MAF) < 0.0001) (Chen et al. 2024b). Model comparison was conducted based on likelihood ratio test (LRT) and Akaike information criterion (AIC) (Supplemental Table S4). By comparing models of different variable combinations, we identified an optimal model, M9, which supports significant effects of three variables (DNVs burden, protein length, and gene age) and the interaction between protein length and DNVs burden (Chi-square test, *p* = 4.59e-66. See details in Supplemental Table S4). The likelihood of being a disease gene increases with four terms: DNVs burden (*D*, coefficient = 0.29; *p* < 2e-16), protein length (*L* on the logarithmic scale, coefficient = 0.22; *p* = 0.00031), gene age (*T*, coefficient = 0.0041; *p* < 2e-16), and the interaction between DNVs burden and the logarithmic protein length (coefficient = -0.027; *p* = 1.27e-9, Supplemental Table S4). The logarithmic scale effect of protein length on the likelihood suggests that normalizing extreme lengths could improve model fit. The negative coefficient for the interaction term between DNVs burden and log-transformed protein length indicates an underlying trade-off between the two factors, suggesting that the impact of mutation burden on the likelihood of a gene being classified as a disease gene decreases as the protein length increases (Supplemental Fig. S2). In other words, while de novo mutation burden generally increases the probability of a gene being a disease gene, this effect is moderated by protein sequence length, with longer proteins showing a diminished impact of mutation burden. The values of variance inflation factor (VIF) for the variables range from 1.03 to 1.55, well below the multicollinearity concern thresholds of 5 to 10, suggesting minimal impact of multicollinearity among these variables on our model.

Consistent with the implications of this model, we found that disease genes tend to be significantly longer than non-disease genes in most branches (Wilcoxon rank sum test, *p* < 0.05), except for br4 and br6 (Supplemental Fig. S3A). The non-significant difference in these two branches may be due to the limited power of comparison caused by the lower sample sizes. We further compared between disease and non-disease genes, using the reported burdens of DNVs (Zhao et al. 2020) and ultra-rare predicted loss-of-function variants (pLOF, minor allele frequency (MAF) < 1 x 10^−5^) from 394,783 exomes (Weiner et al. 2023) (Supplemental Figs. S3B and S3C). For pLOF comparison, we found significant lower burdens of deleterious variants for all branches (Wilcoxon rank sum test, *p* < 0.05). For DNVs comparison, only the primate-specific branch (br6) was not significant, probably due to small sample size. To increase the statistical power of comparison between disease and non-disease genes across different phylostrata, we further collapsed the number of phylostrata to four, starting from the oldest, Euteleostomi, to progressively younger stages of Tetrapoda, Amniota, and Eutheria (Fig. 1A). Based on these four phylostrata, we confirmed that disease genes tend to have significantly longer lengths and higher burdens of deleterious variants than in non-disease genes across all major phylostrata (Wilcoxon rank sum test, *p* < 2.2e-16, Fig. 1D-1E).

### Young genes have lower burdens of DNVs and rare variants than older genes

We further examined the statistical correlation between gene age and burden of potentially deleterious variants. We used datasets from predicted deleterious de novo germline variations (DNVs) (Wang et al. 2022) and genome-wide rare variants from the gnomAD database (Chen et al. 2024b). We found that gene age (mya) positively correlates with DNV burden (Spearman’s rank correlation ρ = 0.181, *p* < 2.2e-16) and rare variant burden (Spearman’s rank correlation ρ = 0.306, *p* < 2.2e-16). When using different methods for human gene age dating—gene-family based (Neme and Tautz 2013) and synteny-based method (Shao et al. 2019)—alongside different datasets on de novo variant (DNVs) burdens (Zhao et al. 2020), we also found a significant correlation between gene age and DNVs burden (Supplemental Fig. S4A and S4B).

New genes could originate from both “copying” and “non-copying” mechanisms (Vonica et al. 2020; Luria et al. 2023; Fleck et al. 2024), with the former from a process of gene duplication and gene traffic (Emerson et al. 2004), while the latter from exclusively sequence evolution (Domazet-Lošo et al. 2007). Thus, genetic novelty of new genes could be either “novelty-by-synteny” or “novelty-by-similarity”. if the novelty resides in the protein sequence itself, then new genes by synteny should have burdens equal to ancient genes that they were copied from. Considering that retrogenes are known genes with “novelty-by-synteny” due to their random insertions, we chose retrogenes and their parental genes to test this possibility. Retrogenes and their parental genes could be easily detected due to their clear structural changes, such as intron loss during retro-transposition and insertion hallmarks (Long and Langley 1993; Miller et al. 2022). We retrieved known pairs of retrogenes and their parental genes from the GenTree database (Supplemental Table S5). We found that over half of retrogenes have lower burden of DNVs and burden score of pLOF variants than their parental genes. In contrast, less than 36% of retrogenes showed higher burden of DNVs and pLOF burden than their parental genes. Thus, this result suggests retrogenes are different from their parental genes in terms of deleterious variant burden, supporting new genes by synteny could lead to functional novelty. Moreover, we compared the primate-specific genes identified only from the synteny-based method and only from the similarity-based method, which are also young genes by synteny and those by similarity, respectively. We found significant higher burdens of DNVs and pLOF in young genes by synteny than those by similarity (Wilcoxon rank sum test, *p* < 0.001). Together, our results suggest that genes of “novelty-by-synteny” are more likely to have disease functions than those of “novelty-by-similarity”, but less likely than their parental genes. These results are consistent with our Supplemental Figs. S4A and S4B that young genes tend to have lower burden of deleterious DNVs than older genes.

### Purifying selection intensifies with gene age and is stronger in disease genes than in non-disease genes

To understand if disease genes evolve under different evolutionary pressures compared to non-disease genes, we compared the Ka/Ks ratio, which is the ratio of the number of nonsynonymous substitutions per nonsynonymous site (Ka) to the number of synonymous substitutions per synonymous site (Ks). Values of Ka/Ks ratios less than 1 suggest a degree of evolutionary constraint (acting against change) (Yang and Bielawski 2000). To ensure similar evolutionary backgrounds, we retrieved the “one-to-one” human-chimpanzee orthologous genes and the corresponding pairwise Ka/Ks ratios (12830 genes) from the Ensembl database (Supplemental Table S6). We also evaluated whether the pattern is consistent with Ka/Ks ratios of human-bonobo and human-macaque orthologs (Supplemental Table S6). Interestingly, Ka/Ks ratios were consistently lower in disease genes than in non-disease genes for human-chimpanzee orthologs (0.250 vs. 0.321), human-bonobo orthologs (0.273 vs. 0.340), and human-macaque orthologs (0.161 vs. 0.213) (Wilcoxon rank sum test, *p* < 2.2e16 for all three datasets). These results revealed that disease genes are under significantly stronger purifying selection than non-disease genes, suggesting an important component of selective pressure in constraining the sequence evolution of disease genes. We observed that Ka/Ks ratios (< 1) increase inversely proportional to gene age, suggesting a trend of relaxed purifying selection on young genes (Figure 1g and Supplemental Figure 2), which is consistent with some previous studies (Carvunis et al. 2012; Prabh et al. 2018; Montañés et al. 2023). For seven age groups, disease genes showed significantly lower Ka/Ks than non-disease genes in five out of seven branches (Supplemental Fig. S5). The non-significant difference in the primate-specific branch (br6) and the therian-specific branch (br4) could be due to lower number of disease genes with Ka/Ks ratios (13 genes in br6 and 56 genes in br4). Notably, despite the relaxation of purifying selection for younger genes, disease genes still tend to show lower Ka/Ks ratios than non-disease genes for gene ages of four phylostrata, suggesting a pattern of stronger purifying selection in disease genes during their macroevolutionary process (Fig. 1G and Supplemental Fig. S5).

We observed a heterogeneous distribution of disease genes underlying 22 HPO-defined anatomical systems, suggesting varied genetic complexity for diseases of different systems (Supplemental Fig. S6A). None of the disease genes was found to impact all 22 systems. In contrast, 6.96% of disease genes (344/4946) were specific to a single system’s abnormality. Notably, four systems – the genitourinary system (with 81 genes), the eyes (68 genes), the ears (63 genes), and the nervous system (55 genes) – collectively represented 77.62% of these system-specific genes (267/344, Supplemental Table S2). The nervous system displayed the highest fraction of diseases genes (79%, Supplemental Fig. S6B). A large proportion of disease genes (93.04%) were linked to abnormalities of at least two systems (4602/4946), indicating that most human disease genes may have a broad phenotypic impact or pleiotropy across multiple anatomical systems. This phenotypic effect across systems might arise from the complex clinical symptoms of rare diseases manifesting in multiple organs, tissues, or systems, potentially indicating levels of pleiotropy (Hodgkin 1998; Paul 2000; Lobo 2008). Compared to commonly used functional inferences based on human gene expression profiles or *in vitro* screening, the comprehensive and deep phenotyping offered by HPO provides a more systematic perspective on the functional roles of human disease genes. Interestingly, we found a moderate but significant negative correlation between the median Ka/Ks ratios and the number of affected anatomical systems in disease genes (linear correlation adjusted R^2^ = 0.38, *p* = 0.0018. Fig. 1H). This implies that disease genes with higher pleiotropy, which impact multiple anatomical systems, face more stringent evolutionary constraints compared to genes with lower pleiotropy (Figure 1H).

### Disease gene emergence rate per million years is similar across macroevolutionary phylostrata

To understand whether different phylostrata have different emergence rates for disease genes, we assessed the disease gene emergence rate per million years across phylostrata from Euteleostomi to Primate (μ). Considering the sampling space variations at different age groups, we calculated μ as the fraction of disease genes per million years at each phylostratum (Fig. 2a). Although the proportions of disease genes were found to gradually increase from young to old phylostrata (Fig. 1a), the rate μ is nearly constant at ∼0.07% per million years for different phylostrata (Fig. 2a). This constant emergence rate of disease genes suggests a continuous and similar fraction of genes evolving to have significant impacts on health during evolutionary history.

**Figure 2.**
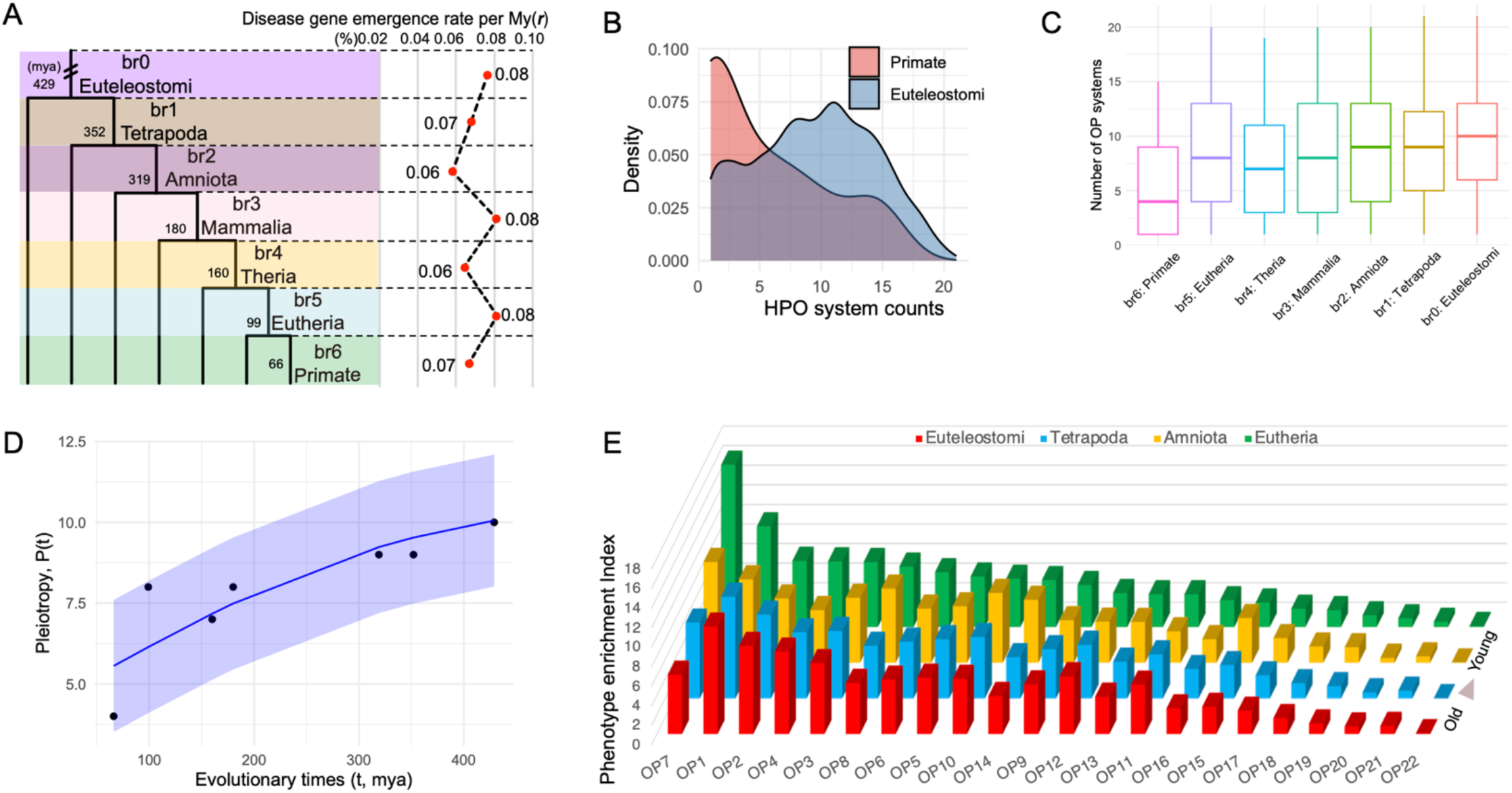
Disease gene emergence rates along phylostrata, OP counts comparison between the youngest and oldest phylostrata, and disease phenotype enrichment index (PEI) along phylostrata. (*A*) The disease-gene emergence rate per million years (*r*) along phylostrata. (*B*) Density distributions showcase numbers of affected organ phenotypic systems (OPs) for genes originated at primate and Euteleostomi phylostrata. (*C*) Boxplot distributions showcase the numbers of affected organ phenotypic systems (OPs) for genes grouped by their phylostrata (median values are 4, 8, 7, 8, 9, 9, 10, from left to right). (*D*) The nonlinear least squares (NLS) regression between pleiotropy score (P) and evolutionary times *t* with the logistic growth function: 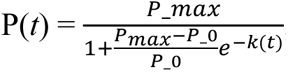, *k* = 1.66, *p* = 0.000787. The 95% confidence interval is shown shade. *P_max* and *P_0* are empirical medians 10 and 4, respectively) (*E*) The distribution of phylostrata and OP for the phenotype enrichment index (PEI). The bar plots, colored differently, represent phylostrata, namely Euteleostomi, Tetrapoda, Amniota, and Eutheria, in ascending order of evolutionary ages. The disease systems (OP) are displayed on the horizontal axis and defined in Fig. 1C. The standard deviations of PEI are 3.67 for Eutheria and approximately 2.79 for older phylostrata.

Using the recently reported average human generation time of 26.9 years (Wang et al. 2023), the most updated number of coding genes (19,831 based on Ensembl v110), and assuming a simplified monogenic model (Richards et al. 2015), we estimated the number of causal genes for rare diseases per individual per generation (μ_d_) as 3.73 x 10^-4^ (= 19,831 x 26.9 x 0.07 x 10^-8^). Using this rate, we can derive the rare disease prevalence rate (r_RD_ = 10,000 x μ_d_), which equates to approximately 4 in 10,000 individuals. This prevalence agrees remarkably well with the European Union definition of rare disease rate prevalence of 5 in 10,000 people (Stolk et al. 2006). This constant emergence rate highlights the idea that young genes continually acquire functions vital for human health, which agrees with the i mportance of young genes in contributing to phenotypic innovations (Kaessmann 2010; Chen et al. 2013; Xia et al. 2021).

### Pleiotropy growth rate is faster in younger genes following a logistic growth pattern

Despite the nearly constant integration of young genes into crucial biological functions (Fig. 2A), it remains uncertain if gene age could influence disease phenotypic spectra (or pleiotropy). The overall distribution of OP (anatomical organ/tissue/system phenotypes) counts for disease genes (Supplemental Fig. S6A) is similar to the distribution of gene expression breadth across tissues (Supplemental Figs. S7A-C). The distribution for OP counts showed that young genes have lower peak and median values than older genes (Figs. 2A-2C). This pattern is consistent with the results that younger genes tend to be expressed in a limited range of tissues, while older genes exhibit a broader expression profile (Supplemental Fig. S7D), which also aligns with previously reported expression profiles (Zhang et al. 2012; Long et al. 2013; Carelli et al. 2016; Miller et al. 2022). We found an increasing trend for median OP numbers from young to old phylostrata (Fig. 2C). Interestingly, the increase rates 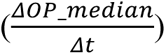 were higher during the younger phylostrata than the older ones (0.12/mya at the Eutheria vs. 0.05/mya at older phylostrata on average, Supplemental Table S7A), suggesting a non-linear and restricted growth model for the level of pleiotropy over time. We applied a logistic growth function and observed a significant pattern: as evolutionary time increases, the level of pleiotropy rises (Fig. 2D, *p* < 0.001). Moreover, the logistic model demonstrates a diminishing marginal growth for pleiotropy over time, indicating that the rate of increase in pleiotropy slows down over evolutionary time. This pattern suggests that while pleiotropy is initially lower in new genes, it increases more rapidly compared to older genes. This result is also consistent with the finding that purifying selection gradually increases over evolutionary time from young to old phylostrata (Figure 1g), which could limit the space of pleiotropy growth in older genes.

### Young genes are highly enriched in the reproductive and nervous system diseases

We found a significant positive correlation between tissue expression breadth and the number of affected disease systems (Spearman’s rank correlation, ρ = 0.138, *p* < 2.2e-16. Supplemental Table S8). To understand the enrichment pattern of disease phenotypes for young and old genes, we introduced a metric of the disease phenotype enrichment index (PEI), which quantifies the range of phenotypes across multiple systems (see Materials and Methods for details). Our findings revealed that the most ancient genes, specifically from Euteleostomi and Tetrapoda, had the strongest PEI association with the nervous system (OP1). Conversely, young genes from Amniota and Eutheria exhibit the highest PEI for disease phenotypes of the genitourinary system (OP7) and the nervous system (OP1), where the genitourinary system (OP7) shows a 38.65% higher PEI than the nervous system (OP1) (Fig. 2E, Supplemental Table S9). Among the 22 disease phenotype systems, only the reproductive system (OP7) showed a steady rise in PEI from older phylostrata to younger ones (Fig. 2E). There were smaller variations in PEI for older phylostrata compared to the more recent Eutheria (∼2.79 vs. 3.67), suggesting that older disease genes impact a greater number of organ systems, as also shown in Fig. 2C. This finding is consistent with the “out-of-testis” hypothesis (Kaessmann 2010), which was built on many observations where the expression patterns of young genes are biased to the testes and can have vital roles in male reproduction. As genes evolve, their expression patterns tend to broaden, potentially leading to phenotypic effects impacting multiple organ systems.

Apart from the reproductive system (OP7), we found that the nervous system (OP1) showed the second highest PEI for Eutherian young disease genes (Fig. 2E). Moreover, 42% of the 19 Primate-specific disease genes with diseases affecting the nervous system (OP1) correlated with phenotypes involving brain size or intellectual development (*CFC1*, *DDX11*, *H4C5*, *NOTCH2NLC*, *NOTCH2NLA*, *NPAP1*, *RRP7A*, and *SMPD4*) (Supplemental Table S2 and Discussion), consistent with the expectation of previous studies based on gene expression (Zhang et al. 2011). Furthermore, young genes emerging during the primate stage are connected to disease phenotypic enrichment in other adaptive systems, particularly in the HPO systems of the head, neck, eyes, and musculoskeletal structure (Fig. 2E). In summary, the primate-specific disease genes could impact phenotypes from both reproductive and non-reproductive systems, particularly the genitourinary, nervous, and musculoskeletal systems (Supplemental Table S2), supporting their roles in both sexual and adaptive evolution.

### Sex chromosomes are enriched for male-reproductive disease genes: the “male X-hemizygosity” effect

Considering the concentration of the youngest disease genes in the reproductive system (Fig. 2E, OP7), we hypothesized that the distribution of disease genes could be skewed across chromosomes. First, we examined the distribution of all disease genes and found a distinct, uneven spread across chromosomes (Fig. 3A and Supplemental Table S10). The X and Y chromosomes contain higher fractions of disease genes compared to autosomes. While autosomes have a linear slope of 0.23 (Fig. 3B, R^2^ = 0.93; *p* = 2.2 x 10^-13^), the proportion of Y chromosomal disease genes is 82.61% higher, at 0.42. Meanwhile, the proportion of X chromosomal disease genes is 30.43% higher than those of autosomes, sitting at 0.301. To understand whether the differences between sex chromosomes and autosomes are related to the reproductive functions, we divided disease genes into reproductive (1285 genes) and non-reproductive (3661 genes) categories based on affected organs (Supplemental Table S10). By fitting the number of disease genes against all genes with gene age information on chromosomes, we observed that the X chromosome exhibited a bias towards reproductive functions. Specifically, on the X chromosome, disease genes affecting non-reproductive systems were slightly fewer than expected (-1.65% excess rate, with 154 observed versus 156.59 expected). The X chromosome displayed a significant surplus of reproductive-related disease genes (observed number 99, expected number 52.73, excess rate 87.75%, *p* < 5.56e-9) (Fig. 3D).

**Figure 3.**
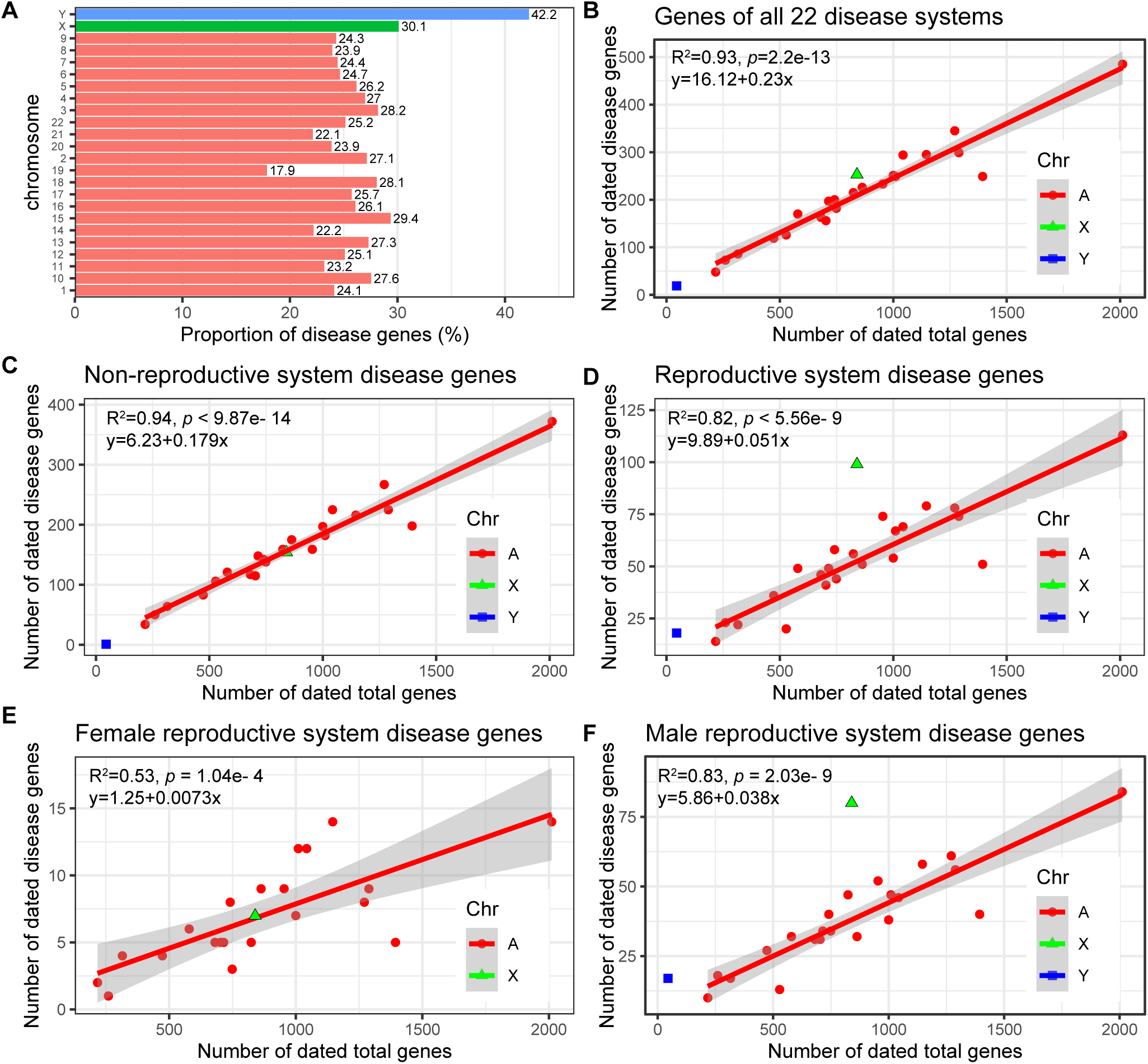
Chromosomal analyses for disease genes. (*A*) The proportions of disease genes across chromosomes. The pink bars represent the autosomes, while green and blue indicate the X and Y chromosomes, respectively. The proportions (%) for different chromosomes are shown above bars. (*B*) The linear regression plotting of disease gene counts against the numbers of total genes with age information on chromosomes. (*C*) Numbers of genes related to the abnormality of non-genitourinary system (non-reproductive system) are plotted against all protein-coding genes on chromosomes with gene age information. (*D*) Numbers of genes related to the abnormality of genitourinary system (the reproductive system) are plotted against all protein-coding genes on chromosomes with gene age information. (*E*) Linear regression of dated disease gene counts against the total numbers of genes on chromosomes for female-specific reproductive disease genes with gene age information. (*F*) Linear regression of disease gene counts against the total numbers of genes on chromosomes for male-specific reproductive disease genes with gene age information. The autosomal linear models are displayed on the top left corner. Note: All linear regression formulas and statistics pertain only to autosomes. “A”, “X”, and “Y” indicate autosomes, X and Y chromosomes, respectively.

Given the sex-imbalanced mode of inheritance for the X chromosome, theoretical model has predicted that purifying selection would remove both dominant female-detrimental mutations and recessive male-detrimental mutations (Rice 1984; Charlesworth et al. 1987). We determined that the ratio of male to female reproductive disease genes (M_disease_/F_disease_ or α) is considerably higher for the X chromosome (80/9 = 8.89) than for autosomes on average (38/21 = 1.81, odds ratio = 16.08, 95% CI: 6.73-38.44, *p* < 0.0001). This suggests a disproportionate representation of disease genes from the male hemizygous X chromosome compared to the female homozygous X. Thus, our analysis indicates that the abundance of disease genes on the X chromosome compared to autosomes is likely due to male-specific functional effects. These results suggest that the overrepresentation of disease genes on the X chromosome primarily results from recessive X-linked inheritance affecting males, rather than dominant effects impacting both sexes.

### Genome-wide excess of male reproductive disease genes: the “faster-X” and “faster-male” effects

To determine which sex (male or female) might influence the biased distribution of reproductive-related genes on different chromosomes, we focused on genes specific to male and female reproductive disease. Based on the HPO terms of abnormalities in reproductive organs and gene age dating, we retrieved 154 female-specific and 945 male-specific disease genes related to the reproductive system with age dating data (Supplemental Table S11). Through linear regression analysis, we assessed the number of sex-specific reproductive disease genes against the total counted genes for each chromosome. We observed strikingly different patterns dependent on sex and chromosomes.

For female reproductive disease genes, the X chromosome followed a linear pattern and did not differ significantly from autosomes (R^2^ = 0.53, *p* = 1.04e-4, Fig. 3E). In contrast, male reproductive disease genes on the X and Y chromosomes showed significant deviations from the autosomes, which followed a linear pattern (R^2^ = 0.82, *p* = 5.56e-9, Fig. 3F). The X chromosome contained 111.75% more male reproductive genes than expected. Moreover, the Y (17/45) and X (80/840) chromosomes had significantly higher ratios of male reproductive disease genes compared to autosomes (averaging 38/853), with odds ratios of 8.48 (95% CI: 4.45–16.17, *p* < 0.0001) and 2.14 (95% CI: 1.44–3.18, *p* = 0.0002), respectively. On the X chromosome, male reproductive genes outnumbered female ones by a factor of 10.43 (80/840 vs. 7/840).

This observation is consistent with the “faster-X hypothesis”, which suggests that purifying selection is more effective at eliminating recessive deleterious mutations on the X chromosome due to the male hemizygosity of the X chromosome (Rice 1984; Charlesworth et al. 1987). Interestingly, a male bias was also evident in reproductive disease genes on autosomes, with the male linear model slope being approximately 4.21 times steeper than that for females (0.038 vs. 0.0073) (Figs. 3E and 3F). Thus, the observed excess of male reproductive disease genes is not solely due to the “faster-X” effect. It might also be influenced by the “faster-male” effect, postulating that the male reproductive system evolves rapidly due to heightened sexual selection pressures on males (Wu and Davis 1993).

### Excess of male reproductive disease genes in younger regions of the X chromosome

While we observed a male bias in reproductive disease genes, the influence of gene ages on this excess remains unclear. We compared gene distribution patterns between older (or ancient, Euteleostomi) and younger (post-Euteleostomi) phylostrata. For female-specific reproductive disease genes, the X chromosome has an excess of ancient genes (25.42%) but a deficiency of young genes (57.16%) (Fig. 4A). Conversely, male-specific reproductive disease genes, younger genes exhibited a higher excess rate than ancient genes (193.96% vs. 80.09%) (Fig. 4A). These patterns suggest an age-dependent functional divergence of genes on the X chromosome, which is consistent with gene expression data (Zhang et al. 2010). The X chromosome is “masculinized” with young, male-biased genes, while old X chromosomal genes tend to be “feminized,” maintaining expression in females (Zhang et al. 2010). On autosomes, the linear regression slope values were higher for male reproductive disease genes than for female ones, both for ancient (0.027 vs. 0.0041) and young genes (0.012 vs. 0.0021) (Fig. 4A). The ratio of male to female reproductive disease gene counts (α) showed a predominantly male-biased trend across phylostrata, with a higher value in the most recent Eutheria (9.75) compared to the ancient phylostrata Euteleostomi and Tetrapoda (6.40 and 3.94, Fig. 4B). A comparison between selection pressure between young and ancient genes revealed no significant difference for female-specific reproductive disease genes, but a significant difference for male-specific ones (Fig. 4C, the Wilcoxon rank-sum test, *p* < 0.0001), indicating that young genes under male-biased sexual selection have fewer evolutionary constraints than older ones (median Ka/Ks ratios 0.35 vs. 0.23).

**Figure 4.**
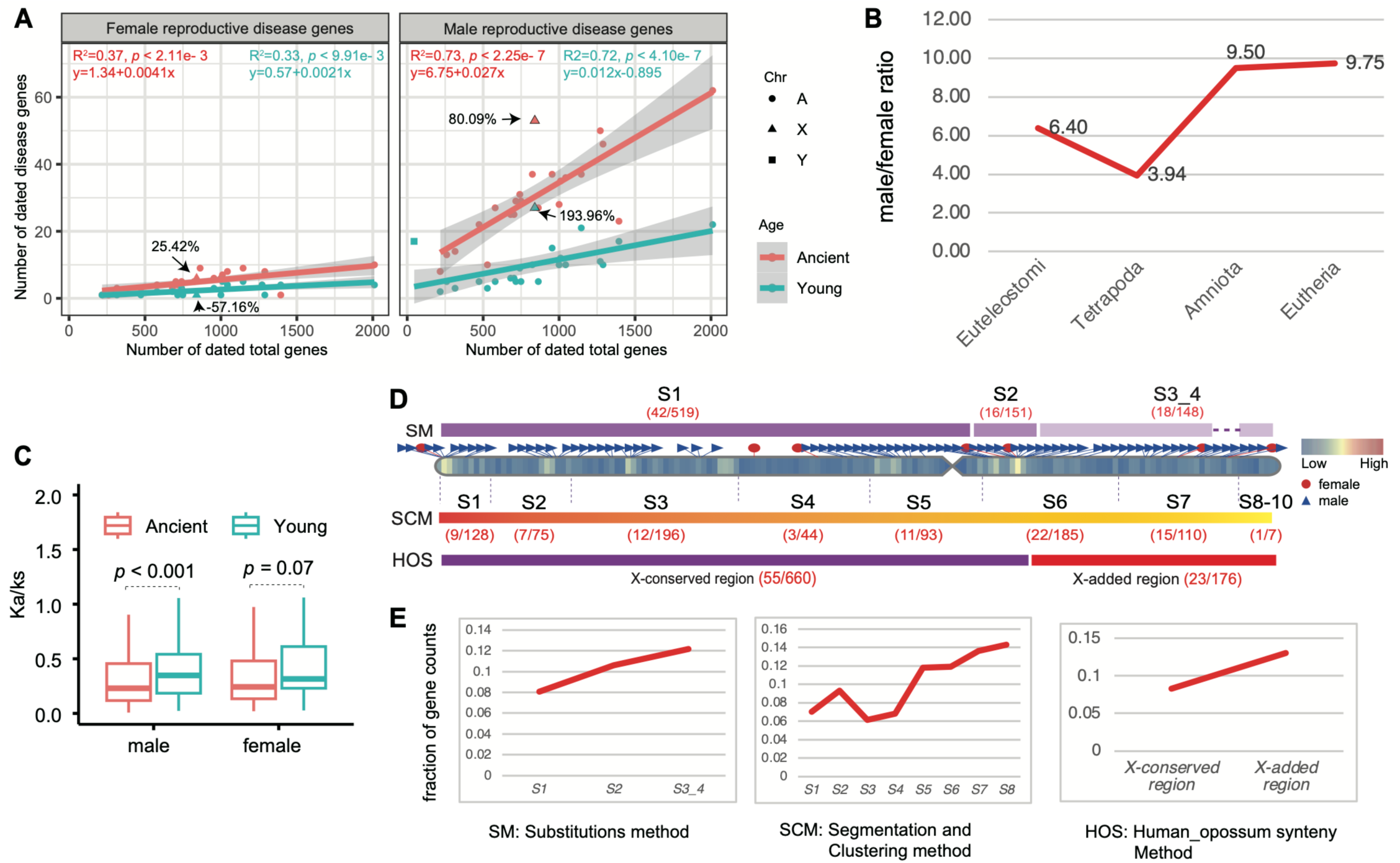
The X chromosomal analyses for disease genes. (*A*) Numbers of female-specific (left) and male-specific reproductive disease genes (right) are plotted against all protein-coding genes with gene ages on chromosomes. The linear formulas fitted for autosomal genes at ancient (Euteleostomi) and younger (post-Euteleostomi) stages are shown in red and blue, respectively. The X chromosome is shown with triangles. (*B*) The ratios of male to female reproductive disease gene numbers (α) across four phylostrata. (C) The comparison of selection pressure (human-chimpanzee pairwise Ka/Ks ratios) for sex-specific reproductive disease genes between the ancient (Euteleostomi or older) and younger (post-Euteleostomi) phylostrata. Only the autosomal comparison is shown, with *p* value from the Wilcoxon test. (*D*) The numbers of male-specific reproductive disease genes (*m*) and the background genes (*b*) within the subregions from old to young in the X chromosome are provided, with the numbers displayed within round brackets for each subregion (*m*/*b*). SM, SCM, and HOS denote three classification methods for X chromosome structure: the substitutions method (SM) (Lahn and Page 1999; McLysaght 2008), the segmentation and clustering method (SCM) (Pandey et al. 2013), and the synteny method (orthologous gene order conservation between human and opossum, HOS) (Ross et al. 2005). (*E*) The fraction of disease genes with male-specific reproductive disease phenotypes within each X chromosomal subregion, as illustrated in (*D*), is presented. The gene coordinates have been updated based on the hg38 reference with liftover. “A”, “X”, and “Y” indicate autosomes, X and Y chromosomes, respectively.

Structurally, the eutherian hemizygous X chromosome comprises an ancestral X-conserved region and a relatively new X-added region (Bellott et al. 2010). The ancestral X-conserved region is shared with the marsupial X chromosome, whereas the X-added region originates from autosomes (Fig. 4D). To understand whether these regions of the human X chromosome might contribute differently to human genetic disease phenotypes, we compared genes within the X-conserved and X-added regions, based on previous studies of evolutionary strata on the X chromosome (Ross et al. 2005; McLysaght 2008; Pandey et al. 2013). After excluding genes in X-PAR (pseudoautosomal regions) regions (Ensembl v110), we found that the proportion of male-specific reproductive disease genes in the X-added region (13.07%, 23/176) exceeds that in the X-conserved region (8.33%, 55/660) (Fig. 4D and 4E, Supplemental Table S12). Moreover, analyses of the evolutionary strata, based on the substitutions method (Lahn and Page 1999) and the segmentation and clustering method (Pandey et al. 2013), consistently showed higher fractions of male-specific reproductive disease genes in younger evolutionary strata than in older ones (Fig. 4E). These observations indicate that, on the X chromosome, young genes could be more susceptible to male-biased sexual selection than old genes, despite their nearly identical hemizygous environment. The higher α in X-linked younger regions could not be attributed to the "male-driven", “faster-X”, “faster-male”, “male X-hemizygosity” effects, as these impact X-linked older and young genes similarly. Instead, the higher α in X-linked young regions may be driven by lower pleiotropy among young genes, allowing novel male-related functions to emerge faster in young genes than in older ones.

## Discussion

### The roles of young genes in human biomedically important phenotypes and innovations

After the discovery of the first disease gene in 1983, which was based on linkage mapping for a Huntington’s disease with a pedigree (Gusella et al. 1983), there has been a rapid advancement in medical genetics research. As of now, this field has identified approximately 20% of human genes (∼4000-5000 genes) associated with rare diseases, "orphan" diseases, and rare forms of common diseases (Asimit et al. 2012; Boycott et al. 2013; Henn et al. 2015; Krumm et al. 2015; Ji et al. 2016; Study 2017; Krausz and Riera-Escamilla 2018; Wen et al. 2018; Almlöf et al. 2019; Povysil et al. 2019; Thuresson et al. 2019; Guo et al. 2021; Oud et al. 2021; Chen et al. 2022b). In our study, we utilized the latest disease gene and clinical phenotype data from HPO annotations (Köhler et al. 2018) and incorporated synteny-based gene age dating to account for new gene duplication events (Shao et al. 2019). Our synteny-based gene age dating reveals that younger genes have lower percentages of disease genes than older genes, qualitatively consistent with previous findings (Domazet-Lošo and Tautz 2008). Recent advances in medical genetics have revealed that Mendelian disorders are predominantly influenced by rare variants due to their significant phenotypic effects (Lee et al. 2014; Guo et al. 2018; Kingdom et al. 2024). However, the correlation between gene age and rare variant burden remains unclear. By analyzing datasets on DNVs and rare variants from large-scale exomic and genomic sequencing aggregated in recent years (Wang et al. 2022; Greene et al. 2023), we found that evolutionary older genes tend to have a higher gene-wise DNVs burden. Logistic regression modeling indicates that protein length, gene age, and DNV burden are positively correlated with the probability of a gene being classified as a disease gene. This suggests that the overrepresentation of disease genes in older evolutionary age groups could result from the combined effect of deleterious variant burden, sequence length, and gene age over evolutionary time under various forms of selection. Despite previous debates on the selective pressures on disease genes (Smith and Eyre-Walker 2003; Domazet-Lošo and Tautz 2008; Chakraborty et al. 2016; Spataro et al. 2017), our comparative analyses of Ka/Ks ratios between humans and primates consistently show stronger purifying selection on disease genes than non-disease genes, indicating evolutionary constraints to remove harmful variants. The phylostratum-wise estimates of the emergence rate of disease genes per million years reveal a steady integration of genes into disease phenotypes, which supports Haldane’s seminal 1937 finding that new deleterious mutations are eliminated at the same rate they occur (Haldane 1937; Keightley 2012).

### Young genes rapidly acquire phenotypes under both sexual and natural selection

The chromosomal distribution of all disease genes shows the excess of disease genes in the X chromosome, which supports the “faster-X effect” (Rice 1984; Charlesworth et al. 1987), that male X-hemizygosity could immediately expose the deleterious X chromosomal mutations to purifying selection. Conversely, the X-chromosome inactivation (XCI) in female cells could alleviate the deleterious phenotypes of disease variants on the X chromosome (Migeon 2020). The X chromosome excess of disease genes is attributed predominantly to that of the male reproductive disease genes. This male-specific bias was not limited to the sex chromosome but also detectable in autosomes. These findings align with the “faster-male” effect, where the reproductive system evolves more rapidly in males than in females due to heightened male-specific sexual selection (Wu and Davis 1993). Intriguingly, of the 22 HPO systems, young genes are enriched in disease phenotypes affecting the reproductive-related system. As genes evolve to be older, there’s a marked decline in both PEI (phenotype enrichment index) and α (the male-to-female ratio of reproductive disease gene numbers). These patterns are consistent with the “out of testis” hypothesis (Kaessmann 2010), which describes the male germline as a birthplace of new genes due to factors including the permissive chromatin state and the immune environment in testis (Vinckenbosch et al. 2006; Bekpen et al. 2018). The “out of testis” hypothesis predicts that genes could gain broader expression patterns and higher phenotypic complexity over evolutionary time (Vinckenbosch et al. 2006). Consistently, we observed a pattern where older sets of disease genes have phenotypes affecting a much broader range of anatomical systems compared to younger genes, which tend to impact fewer systems. The strong enrichment of male reproductive phenotypes for young genes is also consistent with findings from model species that new genes often exhibit male-reproductive expression and functions (Betrán et al. 2002; Heinen et al. 2009), in both Drosophila (Heinen et al. 2009; Gubala et al. 2017; VanKuren and Long 2018) and mammals (Emerson et al. 2004; Jiang et al. 2017). Some new gene duplicates on autosomes are indispensable during male spermatogenesis, to preserve male-specific functions that would otherwise be silenced on the X chromosome due to the meiotic sex chromosome inactivation (MSCI) (Emerson et al. 2004; Zhang et al. 2010; Jiang et al. 2017).

Apart from the reproductive functions, new genes are also enriched for adaptive phenotypes. Previous transcriptomic studies indicate that new genes have excessive upregulation in the human neocortex and under positive selection (Zhang et al. 2011). The brain size enlargement, especially the neocortex expansion over ∼50% the volume of the human brain, ranks among the most extraordinary human phenotypic innovations (Rakic 2009; Zhang et al. 2011). Here, we found that at least 42% of primate-specific disease genes affecting the nervous systems could impact phenotypes related to brain size and intellectual development. For example, *DDX11* is critical in pathology of microcephaly (Pirozzi et al. 2018; Lerner et al. 2020; van Schie et al. 2020; Ma et al. 2022). The *NOTCH2NLA*, *NOTCH2NLB*, and *NOTCH2NLC* may promote human brain size enlargement, due to their functions in neuronal intranuclear inclusion disease (NIID), microcephaly, and macrocephaly (Fiddes et al. 2018; Suzuki et al. 2018; Liu et al. 2022). The *RRP7A* is also a microcephaly disease gene evidenced from patient-derived cells with defects in cell cycle progression and primary cilia resorption (Farooq et al. 2020). The defects of *SMPD4* can lead to a neurodevelopmental disorder characterized by microcephaly and structural brain anomalies (Magini et al. 2019). The *SRGAP2C* accounts for human-specific feature of neoteny and can promote motor and execution skills in mouse and monkey model (Charrier et al. 2012; Dennis et al. 2012; Meng et al. 2023).

New genes were also found to be enriched in other adaptive phenotypes, particularly involving the head and neck, eyes, and musculoskeletal system. Some examples of these primate-specific disease genes include *CFHR3* associated with macular degeneration (Fritsche et al. 2016), *SMPD4* with the retinopathy (Smits et al. 2023), *TUBA3D* with the keratoconus (Hao et al. 2017), *OPN1MW* with loss of color vision (Winderickx et al. 1992; Ueyama et al. 2002), *YY1AP1* with Fibro-muscular dysplasia (Guo et al. 2017), *SMN2* with spinal muscular atrophy (Hahnen et al. 1996), *GH1* with defects in adult bone mass and bone loss (Dennison et al. 2004), *KCNJ18* with thyrotoxicosis complicated by paraplegia and hyporeflexia (Ryan et al. 2010), *TBX5* with the cardiac and limb defects of Holt-Oram syndrome (Basson et al. 1997; Li et al. 1997), and *DUX4* with muscular dystrophy (Lemmers et al. 2012). Additionally, specific functions have also been reported for these young genes. For example, the Y chromosome gene *TBL1Y* could lead to male-specific hearing loss (Di Stazio et al. 2019). Defects in *TUBB8* could lead to complete cleavage failure in fertilized eggs and oocyte maturation arrest (Feng et al. 2016; Yuan et al. 2018; Yao et al. 2022). Interestingly, a previous case study on mice also shows the role of de novo genes in female-specific reproductive functions (Xie et al. 2019). These emerging studies independently support the importance of new genes in phenotypic innovation and sexual selection.

### New genes underlying rapid phenotypic innovations: low pleiotropy as a selective advantage

Our findings raise a question of why new genes can quickly contribute to phenotypic traits that are crucial for both sexual evolution and adaptive innovation. This question could not be fully addressed by previous hypotheses. The "out of testis" theory does not offer specific predictions regarding the propensity of new or young genes to be involved in adaptive traits. Other theories, such as the "male-driven," "faster-X," “male X-hemizygosity”, and "faster-male" theories, cannot explain male-biased functions are more prevalent in young X-linked genes compared to older ones. We found that different phenotypic patterns between young and older genes could be related to differences in their pleiotropy.

The phenomenon of pleiotropy has been recognized or suggested for a considerable time (Mendel 1866; Wright 1984; Barton 1990; Baatz and Wagner 1997). Mendel’s classic paper in 1866 suggests a single factor controls three characters of Pisum (Mendel 1866). Before Mendel, many medical syndromes had already described syndromes characterized by different symptoms and a single “familial” factor (Eckman 1788). In the context of new gene evolution studies, it is established that young genes exhibit higher specificity and narrower expression breadth across tissues (Zhang et al. 2012). In our study, pleiotropy is defined more broadly, involving anatomical systems to understand phenotype evolution (Pyeritz 1989; Lobo 2008; Tyler et al. 2009; Zhang 2023). We reveal a pattern that older genes tend to impact more organs/systems, while young genes display phenotype enrichment in specific organs. Therefore, both phenotype pattern and expression trend across phylostrata suggest that young genes may have lower pleiotropy than older genes.

Numerous theoretical and genomic studies have revealed that pleiotropy impedes evolutionary adaptation, often referred to as the ‘cost of complexity’ (Zeng and Hill 1986; Baatz and Wagner 1997; Orr 2000; Wagner and Zhang 2011; Fraïsse et al. 2018; Quiver and Lachance 2022), while low pleiotropy could foster morphological evolution (Carroll 2005; Wray 2007). The inhibitory effect of pleiotropy on novel adaptation aligns with our observations of the strong purifying selection on genes of high pleiotropy (Wagner and Zhang 2011; Quiver and Lachance 2022) and broad expression patterns (Zhu et al. 2008). As expected, we observed that multi-system and older genes, which exhibit higher pleiotropy, undergo stronger purifying selection. This evolutionary constraint suggests a restricted mutation space to introduce novel traits for old genes due to the “competing interests” of multifunctionality (Fig. 5). The inhibitory pressure could also reduce genetic diversity due to background selection (Charlesworth et al. 1993). The evolution of new genes, especially gene duplicates, serves as a primary mechanism to mitigate pleiotropic effects through subfunctionalization and neofunctionalization (He and Zhang 2005; Guillaume and Otto 2012) and avoid adverse pleiotropy in ancestral copies (Hoekstra and Coyne 2007). The tissue-specific functions of new genes, as a general pattern in numerous organisms, could circumvent adaptive conflicts caused by the multifunctionality of parental genes (Des Marais and Rausher 2008). The reduced pleiotropy in young genes may therefore provide a more diverse mutational space for functional innovation, minimizing unintended pleiotropic trade-offs (Dezső et al. 2008).

**Figure 5.**
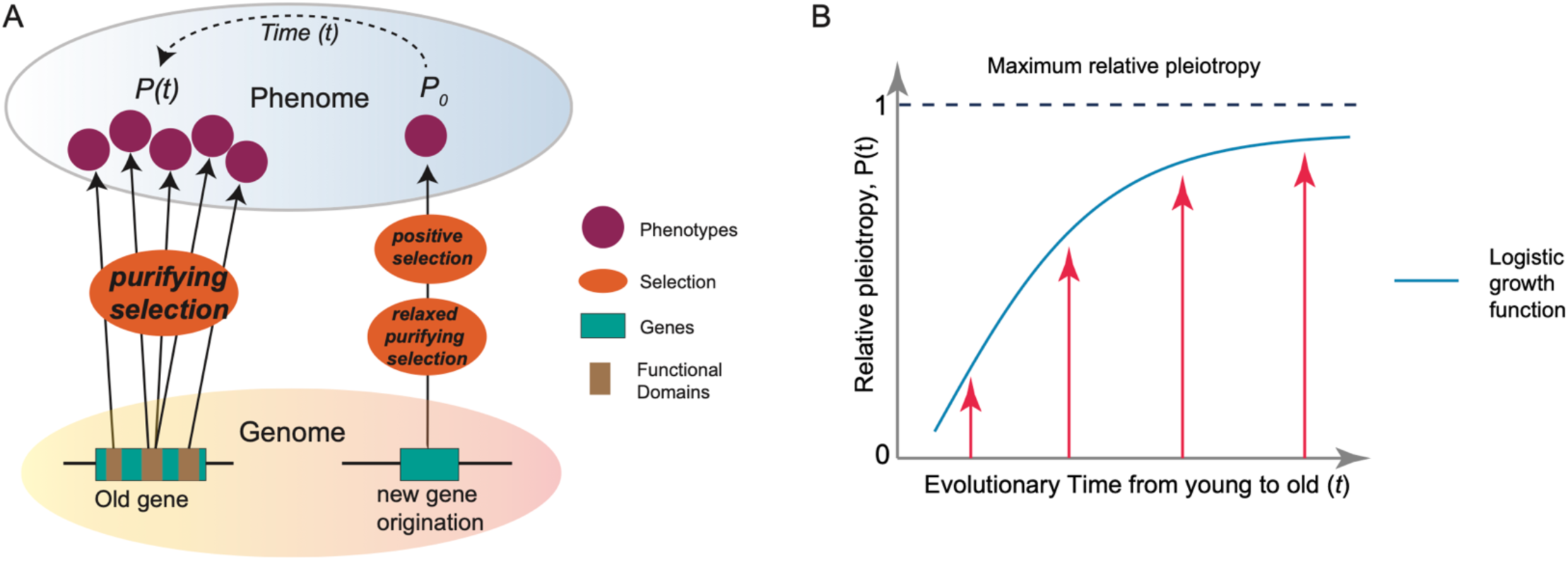
The “pleiotropy-barrier” model. (*A*) The “pleiotropy-barrier” model posits a dynamic process that new genes evolve adaptively more quickly compared to older genes. It suggests that older genes undergo stronger purifying selection because their multiple functions (usually adverse pleiotropy) act as a “barrier-like” factor to hinder fixations of mutations that might otherwise be beneficial for novel phenotypes. (*B*) The logistic function between relative pleiotropy P(*t*) and evolutionary time *t*, 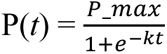, where *P_max* represents the maximum relative pleiotropy. The *k* is the growth rate parameter, which controls how quickly the phenomenon approaches the maximum value. A higher *k* value means faster growth initially.

Here, we propose a “pleiotropy-barrier” hypothesis to explain the relationship between innovation potential and gene ages (Fig. 5A). This model predicts that the capacity for phenotypic innovation is limited by genetic pleiotropy under natural selection, suggesting that genes with lower pleiotropy may have greater potential for functional innovation. Over evolutionary time, the pleiotropy increase follows a logistic growth pattern, where the speed of growth could be higher for younger genes but lower for older genes (Fig. 5B). The multifunctional genes could encounter an escalating “barrier” or “resistance” to the pleiotropy growth. This barrier arises because more functions necessitate stronger selective constraints, which could, in turn, reduce the mutational space for beneficial mutations for novel phenotypes. In contrast, low or absent pleiotropy in new genes allows for a broader and more tunable mutation space under relaxed purifying selection. The permissive environment provides a fertile ground for beneficial mutations to appear with novel functions. Such innovations, initially as polymorphisms within a population, can become advantageous phenotypes and ready responder in certain environment under positive selection. For phenotypes under sexual selection, high pleiotropy may limit the potential to resolve sexual conflict through sex-limited expression (VanKuren et al. 2024). Thus, the “pleiotropy-barrier” model also applies to the evolution of new genes with male-specific functions driven by sexual selection (VanKuren and Long 2018).

The “pleiotropy-barrier” model does not predict a static saturation of pleiotropy in older genes; rather, it emphasizes a continuous and dynamic process between gene age and innovation potential, where gene pleiotropy imposes selective constraints that limit further innovation. This evolving constraint creates a “barrier” that diminishes the potential for genes to acquire new functions, as any beneficial mutations must navigate the complex interplay of existing functions and selective pressures. In contrast, younger genes, which start with low or no pleiotropy, have a greater capacity for evolutionary change. Their low multifunctionality allows them to exploit a wider mutational space, facilitating the development of novel traits and functions. This dynamic process is evident in systems that have undergone significant phenotypic innovations in human evolution, such as the nervous system, musculoskeletal system, and male reproductive system.

Therefore, new or young genes, with lower pleiotropic effect as a selective advantage, not only spur molecular evolution under sexual and natural selection but also, from a medical standpoint, are promising targets for precise medicine, warranting deeper investigation.

## Conclusion

By combining human gene age dating and Mendelian disease phenotyping, we revealed an increasing trend of disease gene proportions over evolutionary time. This growth pattern may be attributed to higher burdens of deleterious variants in older genes. The ratio of health-related genes per million years remains relatively consistent across macroevolutionary phylostrata. Importantly, young genes are preferentially linked to disease phenotypes in the male reproductive system, as well as in systems that have undergone significant phenotypic innovations during primate or human evolution, including the nervous system, head and neck, eyes, and the musculoskeletal system. The enrichment of these disease systems points to the driving forces of both sexual selection and adaptive evolution for young genes. With increasing evolutionary age, genes tend to display fewer specialized functions. Our findings highlight that young genes are likely the frontrunners of molecular evolution for phenotypic innovation. They are actively selected for functional roles driven by adaptive innovation and sexual selection, a process aided by their lower pleiotropy. Therefore, young genes play a pivotal role in addressing a multitude of questions related to the fundamental biology of humans.

## Materials and Methods

### Gene age dating and disease phenotypes

The gene age dating was conducted using an inclusive approach for both autosomes and sex chromosomes (chromosome X and Y). Specifically, for autosomal and X chromosomal genes, we primarily obtained gene ages (phylostrata, branches, or origination stages) from the GenTree database (Zhang et al. 2010; Shao et al. 2019) that is based on Ensembl v95 of human reference genome version hg38 (Flicek et al. 2014). We then trans-mapped the v95 gene list of GenTree into Ensembl gene annotation (v110). The gene age inference in the GenTree database relied on genome-wide synteny and was based on the presence of syntenic blocks obtained from whole-genome alignments between human and outgroup genomes (Zhang et al. 2010; Long et al. 2013; Shao et al. 2019). The most phylogenetically distant branch at which the shared syntenic block was detected marks the latest possible time range when a human gene originated. In comparison to the method based on the similarity of protein families, namely the phylostratigraphic dating (Neme and Tautz 2013), this method employed in GenTree is robust to recent gene duplications (Shao et al. 2019), despite its under-estimation of the number of young genes (Ma et al. 2022). We obtained gene age for human Y genes through the analysis of 15 representative mammals (Cortez et al. 2014). Notably, Y gene ages are defined as the time when they began to evolve independently from their X counter-part or when they translocated from other chromosomes to the Y chromosome due to gene traffic (transposition/translocation) (Cortez et al. 2014). For the remaining Ensembl v110 genes lacking age information, we dated them using the synteny-based method with the gene order information from ENSEMBL database (v110), following the phylogenetic framework of GenTree (Shao et al. 2019). These comprehensive methods resulted in the categorization of 19,665 protein-coding genes into distinct gene age groups, encompassing evolutionary stages from Euteleostomi to the human lineage. The HPO annotation used in this study for phenotypic abnormalities contains disease genes corresponding to 23 major organ/tissue systems (09/19/2023, https://hpo.jax.org/app/data/annotations). This repository synthesizes information from diverse databases, including Orphanet (INSERM 1997; Weinreich et al. 2008), DECIPHER (Wright 2015), and OMIM (Hamosh et al. 2000). After filtering out mitochondrial genes, unplaced genes, RNA genes, and genes related to neoplasm ontology, we obtained gene ages and phenotypic abnormalities (across 22 categories) for 4,946 protein-coding genes. The reproductive system disease genes were retrieved from the ‘phenotype_to_genes.txt’ file by using a grep shell script with the keywords ‘reproduct’, ‘male’, and ‘female’ (neoplasm-related items were removed).

### Logistic regression modeling and model comparison

We retrieved the gene-wise burdens of rare de novo germline variants from multiple studies, including the Gene4Denovo database (68,404 individuals, http://www.genemed.tech/gene4denovo/uploads/gene4denovo/All_De_novo_muta-tions_1.2.txt) (Zhao et al. 2020), and burden scores of ultrarare loss-of-function variants from UK Biobank exomes (394,783 individuals, Supplementary Table 1 in the previous study) (Weiner et al. 2023). The gene-wise burdens of rare variants at the population level were estimated with data from the whole-genome sequencing genotypes and allele frequencies of gnomAD database (Version 4.1.0 of 76,215 individuals, all chromosome VCF files from https://gnomad.broadinstitute.org/downloads) (Wang et al. 2022; Greene et al. 2023). Rare variants were extracted based on MAF lower than 0.0001 across all major human populations (all human male, all human female, African population, non-Finish European population, East Asian population, South Asian population, and Latino/Admixed American population). Other important data, including the files of rare variants and allele frequencies across global human populations (76,215 individuals) and the R script for modeling, could be retrieved from Zenodo (https://zenodo.org/uploads/11000269).

We conducted the stratified Logistic regression to account for effects of multiple predictors and their interactions on the outcome of gene disease states. The disease and non-disease genes were assigned into binary states (“1”, disease genes; “0”, non-disease genes) as response variable. A step-by-step procedure was performed for multiple predictors, which include gene age (*T*, mya), gene length (*L_g_*) or protein length (*L*), DNVs burden (*D*) (Wang et al. 2022), and rare variant burden (*R*) (Chen et al. 2024b). The likelihood ratio test (LRT) and Akaike information criterion (AIC) were used for model comparison. Lower AIC was preferred if the same degree of freedom was detected. The model with significant LRT *p*-value (*p* < 0.05) was chosen when comparing nested models. To account for different scales in variables and potential influence of extreme values, the variables with logarithm treatment were also incorporated in some models. The Variance Inflation Factor (VIF) values were used to account for the multicollinearity among variables. The “glm” package (binomial model) in R platform was used for computing the models (https://www.rdocumentation.org/packages/stats/versions/3.6.2/topics/glm).

### Ka/Ks ratio

Ka/Ks is widely used in evolutionary genetics to estimate the relative strength of purifying selection (Ka/Ks < 1), neutral mutations (Ka/Ks = 1), and potentially beneficial mutations (Ka/Ks > 1) on homologous protein-coding genes. Ka is the number of nonsynonymous substitutions per non-synonymous site, while Ks is the number of synonymous substitutions per synonymous site that is assumed to be neutral. The pairwise Ka/Ks ratios (human-chimpanzee, human-bonobo, and human-macaque) were retrieved from the Ensembl database (v99) (Flicek et al. 2014), as estimated with the Maximum Likelihood algorithm (Yang 2007).

### Disease gene emergence rate per million years (*r*)

To understand the origination tempo of disease genes within different evolutionary phylostrata, we estimated the disease gene emergence rate per million years *r* for disease genes, which is the fractions of disease genes per million years for each evolutionary branch. The calculation is based on the following formula:

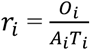

where *r_i_* represents the phenotype integration index for ancestral branch *i*. The *O_i_* indicates the number of disease genes with organ phenotypes in ancestral branch *i*. The denominator *A_i_* is the number of genes with gene age information in branch *i*. The 𝑇_i_ represents the time obtained from the Timetree database (http://www.timetree.org/) (Kumar et al. 2017).

### Pleiotropic modeling with logistic growth function

For each evolutionary phylostratum (*t*), we estimated median OP numbers that genic defects could affect, which serve as the proxy of pleiotropy over evolutionary time (*P*(*t*)) for regression analysis. The logistic growth function was used to fit the correlation with the Nonlinear Least Squares in R.

### Phenotype enrichment along evolutionary stages

The phenotype enrichment along phylostrata was evaluated based on a phenotype enrichment index (PEI). Specifically, within "gene-phenotype" links, there are two types of contributions for a phenotype, which are "one gene, many phenotypes" due to potential pleiotropism as well as "one gene, one phenotype". Considering the weighting differences between these two categories, we estimated the PEI(*i,j*) for a given phenotype (*p_i_*) within an evolutionary stage (*br_j_*) with the following formula.

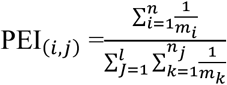

The 𝑚 indicates the number of phenotype(s) one gene can affect, 𝑛 represents the number of genes identified for a given phenotype, and 𝑙 is number of phenotypes within a given evolutionary stage. Considering the genetic complexity of phenotypes, the enrichment index (PEI) firstly adjusted the weights of genes related to a phenotype with the reciprocal value of 𝑚, *i.e*.,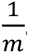. Thus, the more phenotypes a gene affects, the less contributing weight this gene has. Here, *m_i_* is the number of phenotypes affected by the *i*-th gene, n is the total number of genes associated with the specific phenotype *p_i_*, *n_j_* is the number of genes associated with the *j*-th phenotype within the evolutionary stage, and *m_k_* is the number of phenotypes affected by the *k*-th gene within the *j*-th phenotype. Then, we can obtain the accumulative value (*p*) of the adjusted weights of all genes for a specific phenotype within an evolutionary stage. Because of the involvement of multiple phenotypes within an evolutionary stage, we summed weight values for all phenotypes 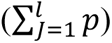 and finally obtained the percentage of each phenotype within each stage 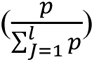 as the enrichment index.

### Linear regression and excessive rate

The linear regression for disease genes and total genes on chromosomes was based on the simple hypothesis that the number of disease genes would be dependent on the number of total genes on chromosomes. The linear regression and statistics were analyzed with R platform. The excessive rate was calculated as the percentages (%) of the vertical difference between a specific data point, which is the number of gene within a chromosome (n), and the expected value based on linear model (n-e) out of the expected value 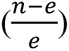.

### Analyses on X-conserved and X-added regions

The Eutherian X chromosome is comprised of the pseudoautosomal regions (PAR), X-conserved region, and X-added region. The regions of two PAR were determined based on NCBI assembly annotation of GRCh38.p13 (X:155701383-156030895 and X:10001-2781479). The X-boundary between X-conserved and X-added regions was determined with Ensembl biomart tool. The "one-to-one" orthologous genes between human and opossum were used for gene synteny identification. The X-conserved region is shared between human and opossum, while X-added region in human has synteny with the autosomal genes of opossum (Ross et al. 2005). The "evolutionary strata" on X were based on previous reports of two methods: substitutions method and the Segmentation and Clustering method (Lahn and Page 1999; McLysaght 2008; Pandey et al. 2013). The coordinates of strata boundaries were up-lifted into hg38 genome with liftover (https://genome.ucsc.edu/cgi-bin/hgLiftOver).

## Acknowledgments

Manyuan Long was supported by the John Simon Guggenheim Memorial Fellowship for Natural Sciences (2022) and the University of Chicago Division of Biological sciences, the National Institutes of Health (1R01GM116113-01A1) and the National Science Foundation (NSF2020667). Deanna Arsala was supported by National Institutes of Health (F32GM146423). We greatly appreciate the constructive discussions with Dr. Stefano Allesina, Dr. Urs C. Schmidt-Ott, and Dr. Greg Dwyer of the University of Chicago, Dr. Anne O’Donnell Luria of the Broad Institute of M.I.T. and Harvard, Dr. Cheng Deng of Western China Hospital, and Dr. Chuanzhu Fan from Wayne State University. Special acknowledgment is given to Xuefei He from Western China Hospital for designing the silhouettes. The authors extend their gratitude to the maintainers and contributors of the HPO data.

## Abbreviations

MAF: minor allele frequency
HPA: the Human Protein Atlas (An expression database)
OP: anatomical organ/tissue/system phenotypes (Fig. 1B)
DNVs: de novo germline mutations/variants
pLOF: predicted loss-of-function variants
HPO: the Human Phenotype Ontology database
LRT: likelihood ratio test
AIC: Akaike information criterion
VIF: variance inflation factor
Ka/Ks: the ratio of the number of nonsynonymous substitutions per nonsynonymous site (Ka) to the number of synonymous substitutions per synonymous site (Ks)
PEI: phenotype enrichment index

