## Supplemental Figure S1 for "Evolutionarily new genes in humans with disease phenotypes reveal functional enrichment patterns shaped by adaptive innovation and sexual selection"

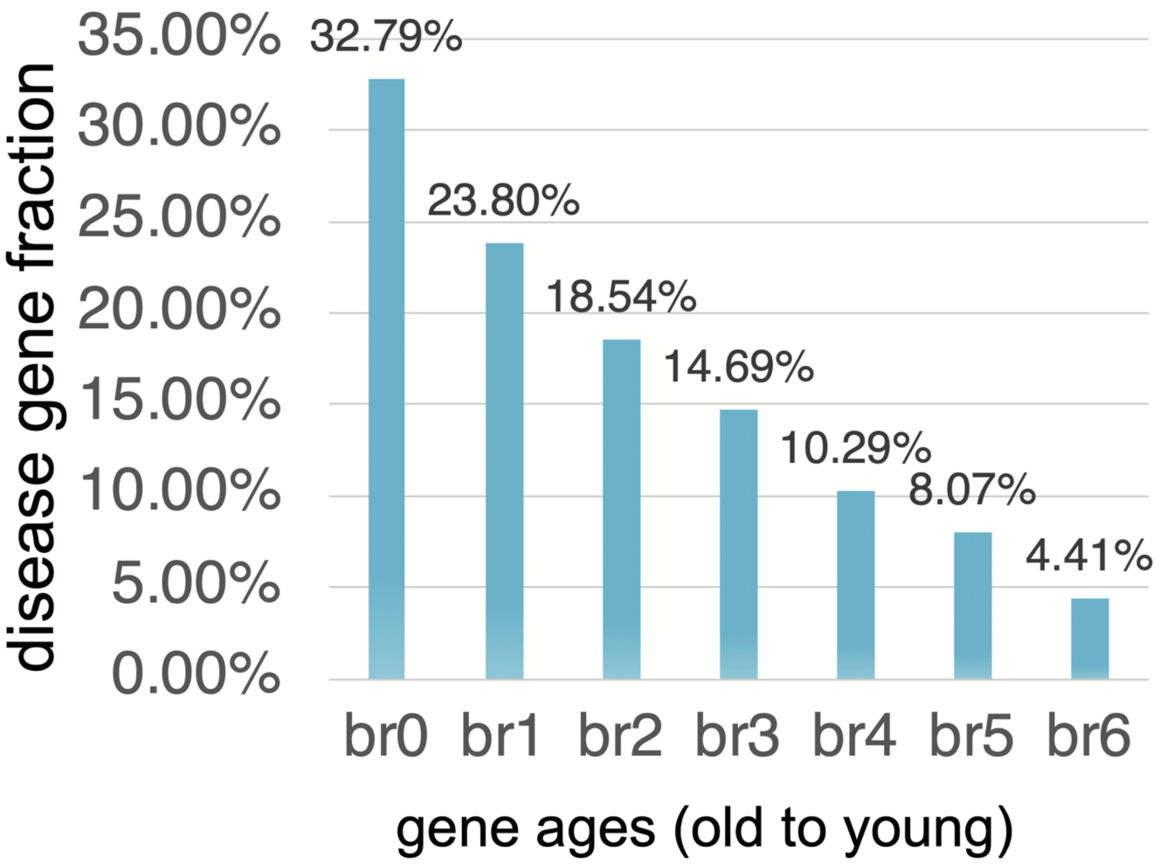


The fractions of disease genes for seven age groups (phylostrata). The horizontal axis shows the seven age groups and the vertical one shows the fractions of disease genes out of each age group. The “br” indicates “branch”, which is also age group or phylostratum.
