## Supplemental Figure S2 for "Evolutionarily new genes in humans with disease phenotypes reveal functional enrichment patterns shaped by adaptive innovation and sexual selection"

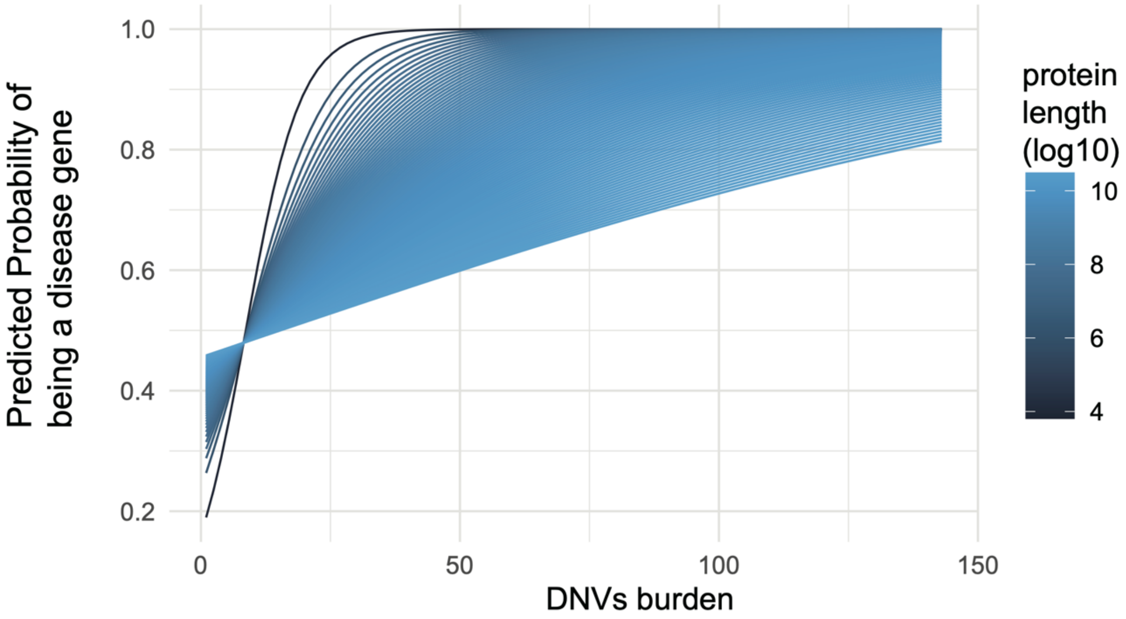


The interaction plot between DNVs burden and protein length (at logarithm scale) for the predicted probability of being a disease gene. The details of full model are shown in Supplemental Table 4.
