## Supplemental Figure S3 for "Evolutionarily new genes in humans with disease phenotypes reveal functional enrichment patterns shaped by adaptive innovation and sexual selection"

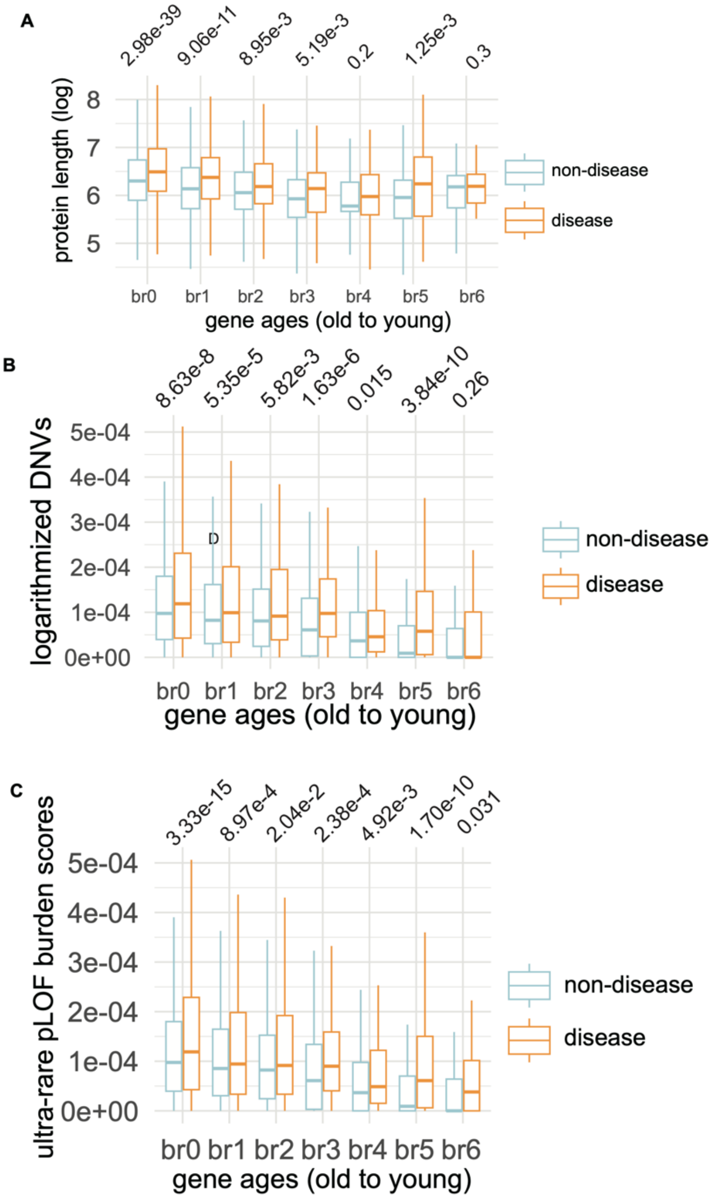


The relationship between multiple features (protein length, burden of DNVs, ultra-rare pLOF burden score) and seven gene age groups (phylostrata). (*A*) The comparison of protein lengths across gene ages between disease genes and non-disease genes. (*B*) The comparison of DNVs burdens across gene ages between disease genes and non-disease genes. (*C*) The comparison of ultra-rare pLOF burden scores across gene ages between disease genes and non-disease genes.
