## Supplemental Figure S4 for "Evolutionarily new genes in humans with disease phenotypes reveal functional enrichment patterns shaped by adaptive innovation and sexual selection"

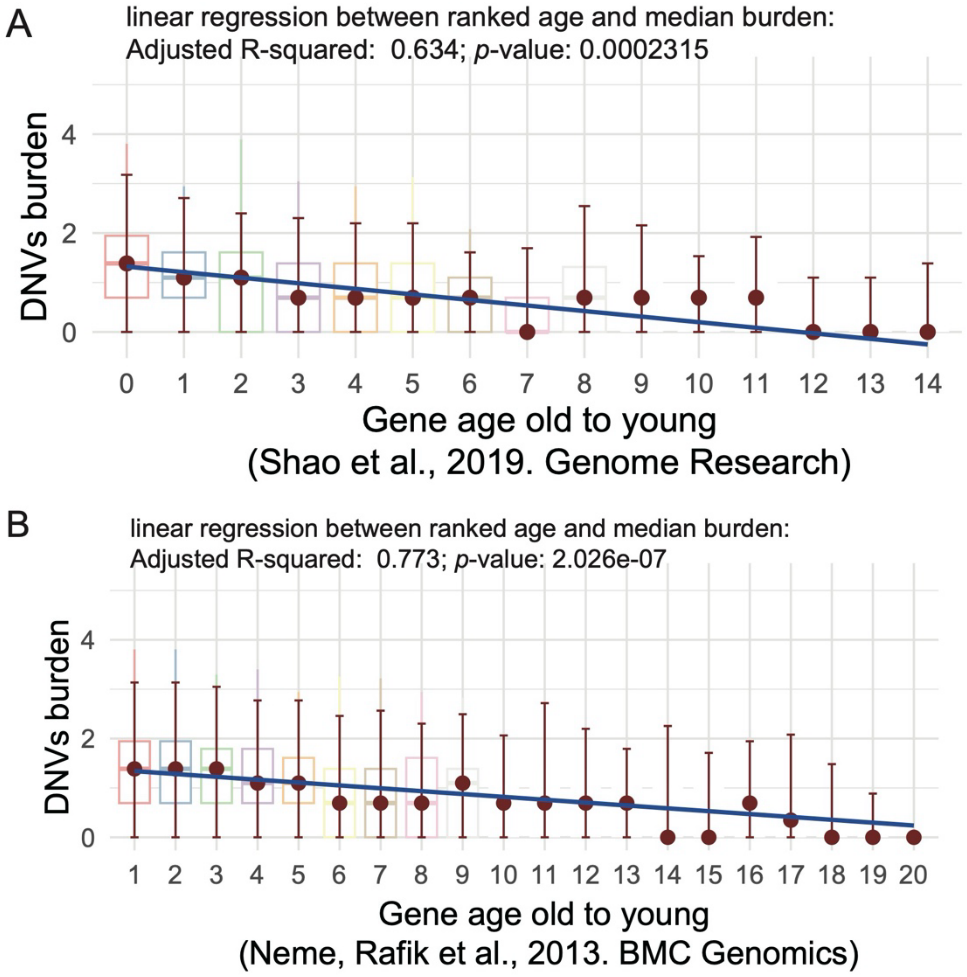


The relationship between two types of gene age dating and the DNVs burden. (*A*)The relationship between gene-wise DNVs burden from 68,404 individuals (Zhao et al. 2020) and the previously reported synteny-based gene age (Shao et al. 2019). (*B*) The relationship between gene-wise DNVs burden from 68,404 individuals (Zhao et al. 2020)and gene-family based gene age (Neme and Tautz 2013). Note: the linear models are between median values and ranked ages. The significance *p* values are shown above age groups (the one-tail Wilcoxon rank sum test with continuity correction).
