## Supplemental Figure S5 for "Evolutionarily new genes in humans with disease phenotypes reveal functional enrichment patterns shaped by adaptive innovation and sexual selection"

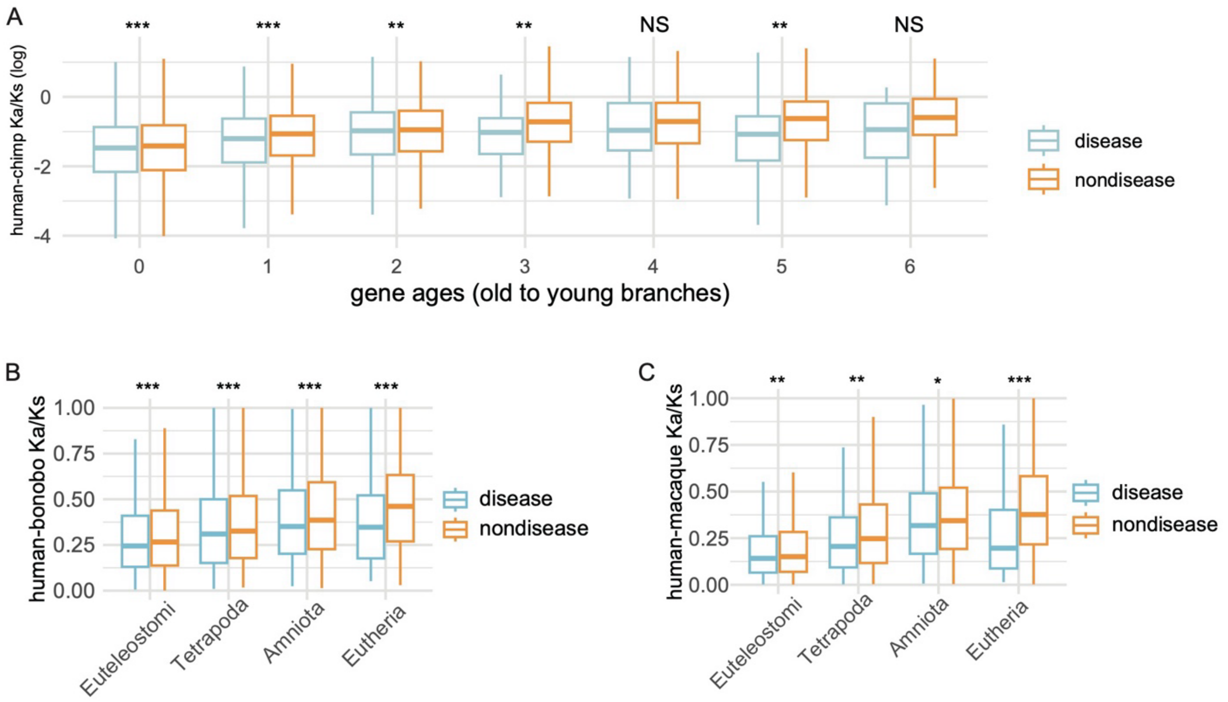


The pairwise Ka/Ks ratios from the Ensembl database based on the Maximum Likelihood estimation for “one-to-one” orthologs between human and other species. (*A*) The pairwise Ka/Ks ratios between human and chimpanzee across seven age groups. (*B*) The pairwise Ka/Ks ratios between human and bonobo across four age groups. (*C*) The pairwise Ka/Ks ratios between human and macaque across four age groups. Only genes under purifying selection are visualized (Ka/Ks < 1). Note: significance levels are based on the Wilcoxon rank sum test comparing disease genes and non-disease genes (one tail test). ”*”, ”**”, “***” indicate $p < 0.05$, $< 0.01$, $< 0.001$, respectively.
