## Supplemental Figure S6 for "Evolutionarily new genes in humans with disease phenotypes reveal functional enrichment patterns shaped by adaptive innovation and sexual selection"

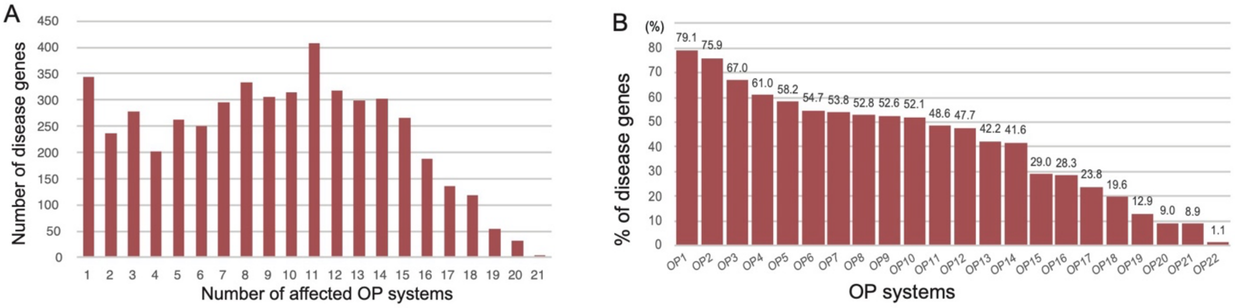


The numbers of disease genes affecting single disease system (OP count = 1) and multiple OP systems (two or more OPs). (*A*) The distribution disease gene counts along the OP numbers. The horizontal axis is based on the numbers of disease systems that a gene defect could impact, and the vertical axis is the number of disease genes. (*B*) The percentage of disease genes for each disease system (OP1 to OP22). The percentage is based on the number of disease genes for certain OP out of all disease genes. The definition of OP systems is consistent with Fig. 1B.
