## Supplemental Figure S7 for "Evolutionarily new genes in humans with disease phenotypes reveal functional enrichment patterns shaped by adaptive innovation and sexual selection"

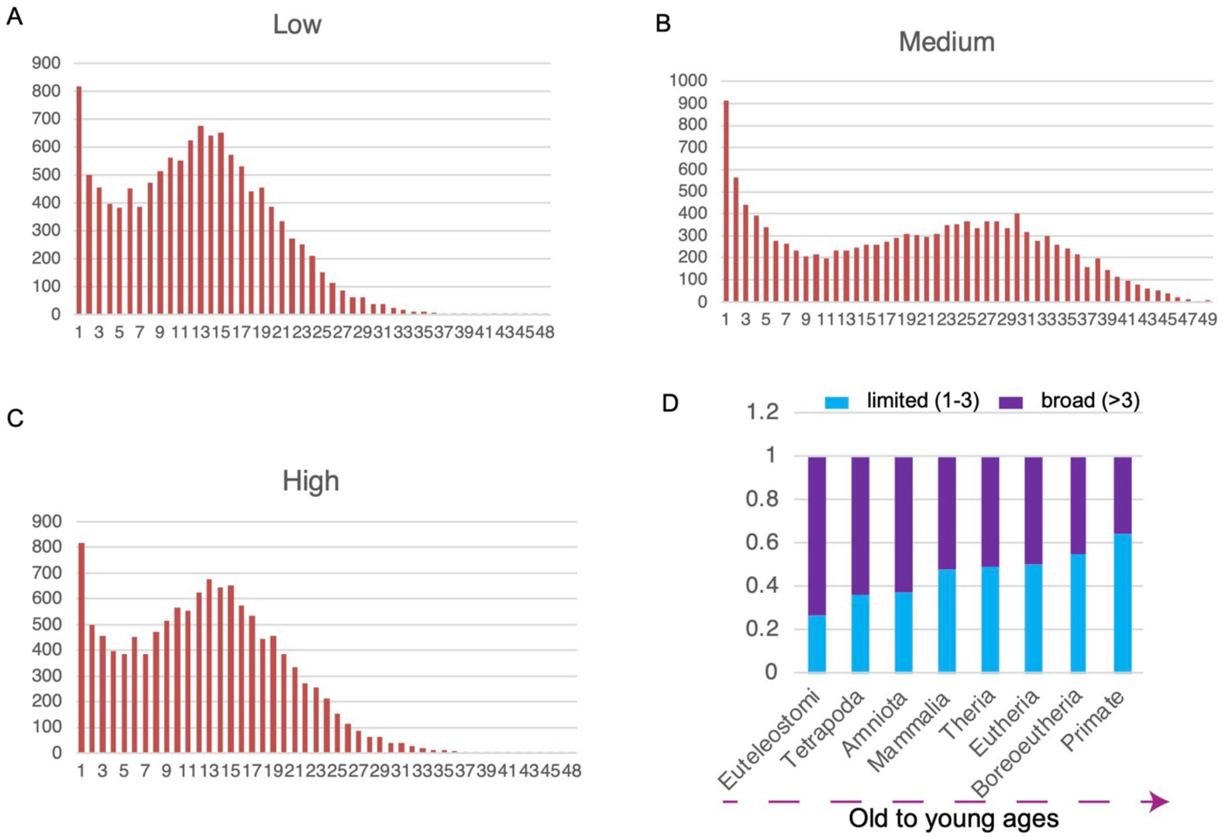


The distribution of gene expression breadth (the number of tissues) with RNASEQ data (The Human Protein Atlas, or HPA, normal tissues). (*A*) The distribution of counts of normal tissues with low levels of gene expression (based on HPA annotation). (*B*) The distribution of counts of normal tissues with medium levels of gene expression (based on HPA annotation). (*C*) The distribution of counts of normal tissues with high levels of gene expression (HPA annotation). (*D*) The distribution percentages of genes expressed in multiple tissues (>3) compared to those expressed in a limited number of tissues (<=3) across different evolutionary ages (only "HIGH" expression genes based on HPA are used).
